# Forecasting the potential distribution of the invasive vegetable leafminer using ‘top-down’ and ‘bottom-up’ models

**DOI:** 10.1101/866996

**Authors:** James L. Maino, Elia I. Pirtle, Peter M. Ridland, Paul A. Umina

## Abstract

The vegetable leafminer, *Liriomyza sativae*, is an internationally significant pest of vegetable and flower crops, that was detected for the first time on the Australian mainland in 2015. Due to the early stage of its invasion in Australia, it is unclear how climatic conditions are likely to support and potentially restrict the distribution of *L. sativae* as it expands into a novel range and threatens agricultural production regions. Here we predicted the future establishment potential of *L. sativae* in Australia, using both a novel ‘bottom-up’ process-based model and a popular ‘top-down’ correlative species distribution model (SDM), leveraging the unique strengths of each approach. Newly compiled global distribution data spanning 42 countries was used to validate the process-based model of establishment potential based on intrinsic population growth rates, as well as parameterise the correlative SDM. Both modelling approaches successfully captured the international distribution of *L. sativae* based on environmental variables and predicted the high suitability of non-occupied ranges, including northern regions of Australia. The largely unfilled climatic niche available to *L. sativa*e in Australia demonstrates the early stage of its Australian invasion, and highlights locations where important vegetable and nursery production regions in Australia are highly vulnerable to *L. sativae* establishment.

## 1. Introduction

There is a growing appreciation of the biosecurity risks posed by the increasing domestic and international flow of goods and people as economies increase in scale and connectivity (Wichmann et al. 2009). Prioritization of high priority areas can be aided by species distribution models (SDMs) which use environmental data to explain spatial patterns in the occurrence likelihood of species (Kearney and Porter 2009; Elith and Leathwick 2009). Species distribution models (SDMs) based on statistical inference or ‘top-down’ inference from occupied/unoccupied locations and associated environmental conditions are frequently used to estimate environmental suitability for species of interest (Phillips et al. 2006; Elith and Leathwick 2009). Such approaches are useful for identifying complex, and often latent and interacting, ecological processes, so long as appropriate environmental covariates that sufficiently characterise the species’ niche can be selected (Phillips et al. 2006; Guillera-Arroita et al. 2015). However, such approaches are challenged by the inherent risk of extrapolating statistical inference to novel conditions, such as spatial ranges not yet invaded (e.g. Dormann et al. 2012; Maino et al. 2016). These problems are exacerbated when using biased occurrence data (e.g. more reports in highly populated areas) or less proximate environmental variables that do not have a constant relationship through space with the limiting process (e.g. using altitude instead of minimum winter temperature when cold tolerance is limiting) (Austin 2007). One solution is to incorporate additional biological and ecological knowledge of the underlying processes that restrict a species’ distribution, such as resource requirements or physiological limits, to make predictions from the ‘bottom-up’ (in contrast to ‘top-down’ inference from occurrence data) (Kearney and Porter 2009). However, such process-based models rely on the mechanistic understanding of the scientist, who may not be able to detect complex and high dimensional interactions between environmental processes that are easily identified through statistical approaches. The comparative strengths of these two approaches can offer mutual benefit. When physiological knowledge of the limiting processes are incomplete, parameters can be inferred in much the same way as top-down SDMs; the key difference being the physical dimensions, theoretical coherence, and interpretability of the fitted parameters, in contrast to risk indices produced from summing weighted quantities of mixed dimensions.

Within Australia, a list of 40 ‘high risk’ biosecurity species has recently been identified by government (Department of Agriculture and Water Resources 2016), which includes several species of Agromyzidae flies, including the vegetable leafminer (*Liriomyza sativae*), the American serpentine leafminer (*Liriomyza trifolii*) the pea leafminer (*Liriomyza huidobrensis*) and the tomato leafminer (*Liriomyza bryoniae).* These species have each been identified as high risk to Australian agriculture (Jovicich 2009). In 2008, *L. sativae* was detected for the first time throughout the north Australian islands of Torres Strait (Blacket et al. 2015), and then on the Australian mainland at Seisia in 2015 (IPPC 2017). The pest has not yet been detected in any other regions of Australia despite ongoing surveillance efforts. As a highly polyphagous insect capable of infesting a broad range of vegetable and flower crops, *L. sativae* represents a significant threat to the production value of vegetables (valued at $ 3.9b in 2016/17) and nurseries (valued at $ 1.6b in 2016/17) (ABS 2018).

The majority of *L. sativae* damage occurs during larval feeding between the upper and lower leaf surface, which curtails photosynthetic ability and reduces marketability of some crops (Parrella 1987). *L. sativae* is also prone to evolving insecticide resistance, making control and eradication difficult (Mason et al. 1987). Overseas, where *L. sativae* is established in production areas, insecticide-based control disrupts beneficial predators and parasitoids, leading to secondary outbreaks of *L. sativae* (Parrella 1987). The risk of inappropriate chemical applications is particularly high during incursions into new regions due to unfamiliarity with the pest. Before *L. sativae* reaches agricultural areas, preparedness can be enhanced through awareness and response plans. Preparedness is not only paramount to effective pest management, but also helps to reduce subsequent rates of spread and establishment. However, due to limited resources for awareness raising and preparedness for exotic pests, high risk regions must be prioritised based on the likelihood *L. sativae* will be able to establish and grow to numbers large enough to cause crop destruction and facilitate further spread.

Here, in a comparison of ‘top-down’ and ‘bottom-up’ approaches, we forecast the establishment potential of *L. sativae* to: (i) identify environmental drivers of the known current global distribution of *L. sativae*; (ii) identify areas in Australia of high climatic suitability for *L. sativae*; (iii) compare predicted areas of suitability in two opposing SDM approaches and; (iv) identity vegetable production areas at highest vulnerability to *L. sativae* as measured by climatic suitability.

## 2. Methods

To estimate the potential distribution of *L. sativae* in Australia based on population growth potential we: (i) compiled empirical data on the temperature response of intrinsic population growth, (ii) compiled ecophysiological data on the mortality response to extreme environmental stressors, including temperature and moisture stress, (iii) projected estimated population growth across the world via modern global climatic data sets, (iv) validated the predicted distributions using available occurrence data, and (v) compared results from this approach with a widely applied ‘top-down’ SDM approach using the MaxEnt algorithm.

### Population growth potential

The intrinsic rate of population growth is the exponential growth rate of a population when growth is not limited by any density dependent factors. More formally, if the change in population size *N* with time *t* is expressed as 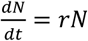, then *r* is the intrinsic population growth rate (individuals per day per individual), which can be decomposed into per capita reproduction and mortality rate. The intrinsic growth rate parameter *r* depends strongly on temperature, with population growth inhibited at low and high temperatures (Haghani et al. 2006), and can be described using a variety of non-linear functions. Here, on the basis that negative population growth cannot be reliably measured under common population cohort studies where reproduction must be positive (i.e. a negatively growing population cannot be maintained in the laboratory), we separated the parameterisation of the positive intrinsic growth rate *r*_*p*_ from negative growth *r*_*n*_. The temperature response of positive growth rate was modelled using a formulation of the Sharpe and DeMichele model (Schoolfield et al. 1981) and parameterised from empirical data (Zhang et al. 2000; Haghani et al. 2006; Chien and Chang 2007) and non-linear least squares regression (Figure 1).

**Figure 1.**
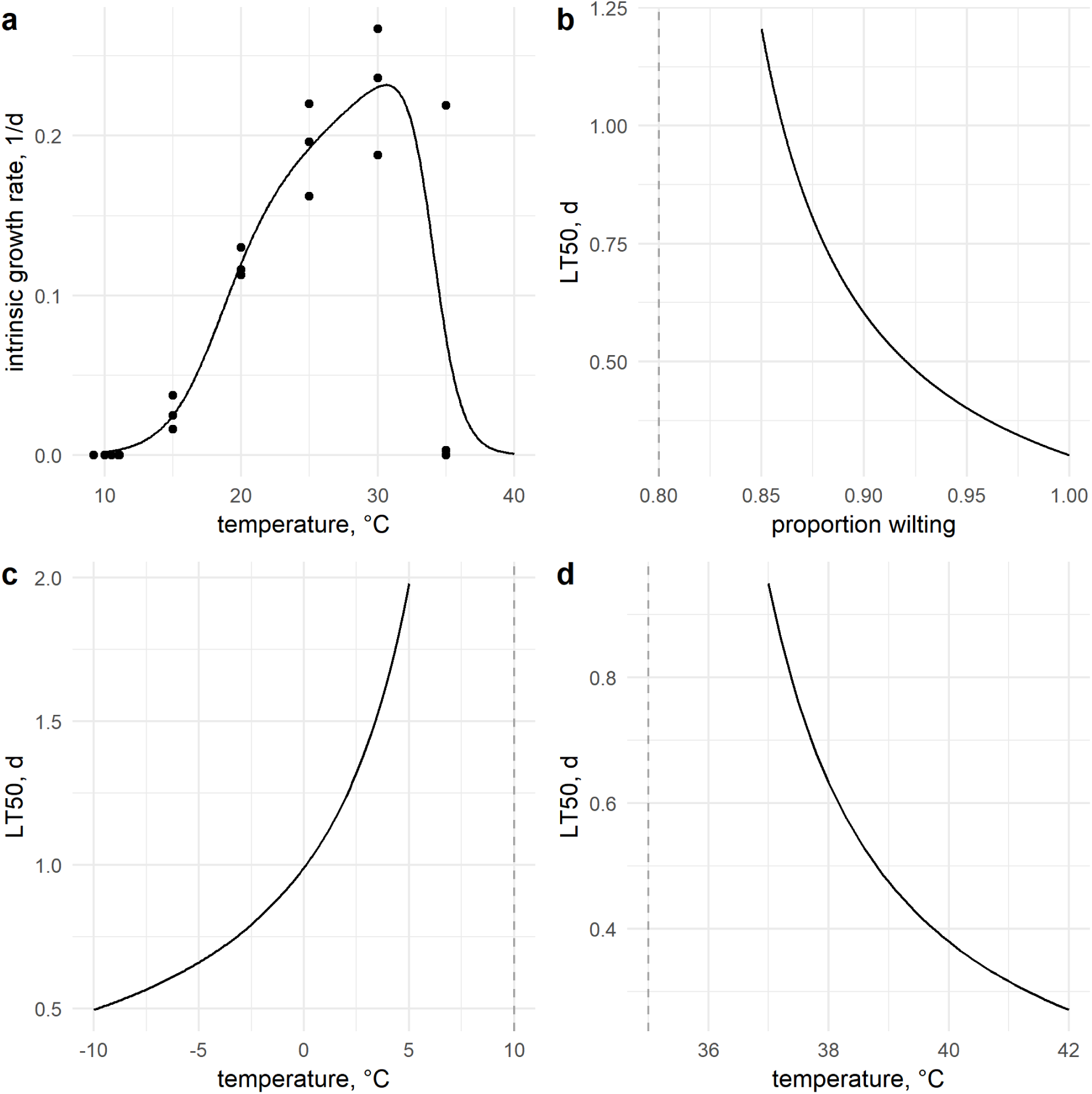
Estimated *Liriomyza sativae* intrinsic population growth rate based on measured empirical life table data (Zhang et al. 2000; Haghani et al. 2006; Chien and Chang 2007) (**a**), and stressor mortality rates expressed as time to 50% mortality (LT_50_) from desiccation (**b**), cold (**c**) and heat (**d**). Dashed lines indicate the critical threshold. See Table 1 for justification of the stress responses.

### Population mortality from extreme conditions

Extreme stressor mortality can be assumed to occur once an environmental variable *s* exceeds some threshold (e.g. critical thermal maximum), beyond which the mortality rate scales approximately linearly with the depth of the stressor (Enriquez and Colinet 2017). Stressor induced mortality was incorporated through quantifying the threshold *s*_*c*_ beyond which stress associated mortality commences, and the mortality rate parameter *m*_*s*_ which reflects the per capita mortality per stress unit per time (e.g. degrees beyond the stress threshold per day). The mortality rate for each stressor *s* was incorporated as 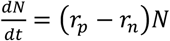 where *r*_*n*_ = ∑_*s*_ *f*(*s, c*_*s*_)*m*_*s*_ and *f*(*s, c*_*s*_) is a function that provides the positive units by which *s* exceeds *s*_*c*_. For example, 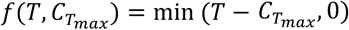 when *T* is temperature 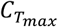 is the heat stress threshold.

**Table 1.** Parameters for critical thresholds and mortality rates for key environmental stressors for *Liriomyza sativae*.

| Parameter | Description | Value | Justification |
| --- | --- | --- | --- |
| $C_{T_{min}}$ | Critical minimum temperature, °C | 10 | The lower developmental threshold has been estimated in several studies as 10.7°C (Tokumaru and Abe 2003), 11.1°C (Sakamaki et al. 2003), and 9.6°C (Zeng et al. 1998). |
| $m_{T_{min}}$ | Mortality rate per cold stress, °C/d | -0.07 | Adult survival after 24h was ~30% at -6.3°C (Oatman and Michelbacher 1959), which we use to solve for $m_{T_{min}} = \frac{\ln(0.30)}{C_{T_{min}} - (-6.3)}$. |
| $C_{T_{max}}$ | Critical maximum temperature, °C | 35 | The upper developmental threshold has been estimated in several studies as 35°C (Oatman and Michelbacher 1959; Chen et al. 1999; Haghani et al. 2006; Chien and Chang 2007), however Zhang et al. (2000) found some population persistence. |
| $m_{T_{max}}$ | Mortality rate per heat stress, °C/d | 0.365 | Mortality rates at extreme heat are unavailable for *L. sativae*, however in the related species *L. bryoniae*, there was low recovery when adults were brought to 41.3°C (Hands 2013), which we take as 10% survival to parameterise $m_{T_{max}} = \frac{\ln(0.1)}{C_{T_{max}} - 41.3}$. |
| $C_{wilting}$ | Critical wilting fraction, - | 0.80 | There is little data on *L. sativae* under moisture stress, and so 80% of plants wilting is arbitrarily assumed as the moisture stress threshold. |
| $m_{wilting}$ | Mortality rate per desiccation stress, | 11.5 | At a soil moisture of 0%, only 10 percent of pupa had successfully completed pupation (Zhang et al. 2004), which |
| | 1/d | | we use to parameterise $m_{\text{wilting}} = \frac{\ln(0.1)}{c_{\text{wilting}} - 1}$. |

This relied on the simplifying assumption that different stressors contribute additively to mortality rate, but conveniently allowed the mortality response of stressors to be partitioned and calculated separately to intrinsic growth rate, which aids parameterisation. More complicated stress responses are more likely to capture reality more completely, but as demonstrated below, the availability of data on detailed mortality responses to climatic stressors for *L. sativae* is lacking.

Here we considered the thermal stressors (critical maxima and minima) as well as moisture stress. *L. sativae* is a herbivorous species, so rather than soil moisture we used the proportion of plants at wilting point to be more relevant, which considers the effects of soil type on water potential. Once these thresholds have been exceeded, the mortality rate *m*_*s*_ for each stressor *s* was estimated from previous studies using the solution to the differential equation when intrinsic growth rate is negative, 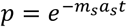 where *p* is the surviving proportion and *a*_*s*_ is the accumulated stress units until time *t*. Table 1 provides estimates for threshold parameters for climatic stressors.

### Simulating global population growth potential

The Soil Moisture Active Passive (SMAP) data products derived from the SMAP satellite mission were used to estimate various climatic conditions relevant to habitat suitability for *L. sativae* (Entekhabi et al. 2010). We used SMAP Level 4 data products, which are model-derived value-added products that combine SMAP satellite observation with a land surface model and observations-based meteorological forcing data, including precipitation and temperature to provide global gridded climatic and environmental data at the 9km resolution every 3 hours from April 2015 since satellite mission commencement (Reichle et al. 2017). Validation analysis of SMAP has revealed that SMAP data products are generally better than other comparable data sets (Reichle et al. 2017). Three SMAP data fields were used to define climatic stressors included ‘surface_temp’ (the mean land surface temperature (K)), and ‘land_fraction_wilting’ (the fractional land area that is at wilting point based on soil moisture at 0-5 cm (m^3^ m^-3^) and soil type).

Using averaged daily SMAP climatic data and the parameters defined in Table 1, for each month the mean daily stress limited growth rate *r -* ∑_*s*_ *f*(*s, s*_*c*_)*m*_*s*_ was calculated using 3-hourly timesteps. These monthly values were then used to derive summary statistics, such as mean positive monthly population growth, or number of months with positive population growth per year. These summaries enabled us to collapse the seasonal variation in growth potential into one temporal dimension to aid visualisation. We summarised these temporal fluctuations by summing the positive growth across months to estimate maximum growth potential, as well as mean growth rate, which includes negative growth and thus represents population persistence.

### MaxEnt model

In addition to the ‘bottom-up’ model described above, we applied the commonly used ‘top-down’ SDM approach using the software Maximum Entropy Modelling v.3.3.3 (Phillips et al. 2006), which has been found to perform well on presence-only data sets (González-Irusta et al. 2015). As environmental covariates, we used BioClim variables (version 2) from the WorldClim database (Fick and Hijmans 2017), which includes global monthly climatic data of minimum, maximum and means for both temperature and rainfall at the 5-minute resolution. To reduce collinearity of the covariates and increase parsimony of the model we used a lowly-correlated subset of the BioClim variables selected for Dipteran horticultural pests (Ørsted and Ørsted 2019), which included ‘annual mean temperature’ (BIO 1), ‘mean diurnal range’ (BIO 2), ‘temperature seasonality (CV)’ (BIO 4), ‘maximum temperature of the warmest month’ (BIO 5), ‘minimum temperature of the coldest month’ (BIO 6), ‘temperature annual range’ (BIO 7), ‘mean temperature of coldest quarter’ (BIO 11), ‘annual precipitation’ (BIO 12), and ‘precipitation of the driest quarter’ (BIO 17).

To reduce sampling bias in the occurrence data set, only a single record for each occupied grid cell was used, with pseudo-absences selected from a 250 km radius around each occupied grid cell, which was assumed to have been surveyed (Merow et al. 2013). In a 5-fold cross-validation fitting procedure, 10,000 random background samples were used as pseudo-absence points. Default settings for features construction and regularization were used. Habitat suitability is shown on the probability scale ranging from 0 to 1, representing unsuitable to optimal habitat.

### Comparison of production regions

Seasonal and regional variation in estimated population growth rates was explored for five key growing regions selected based on production volume of affected agricultural commodities and variation in climatic conditions. This included Lakeland, Bundaberg, Kununurra, Werribee, Mildura (locations indicated in the map in Figure 5). Seisia was also included as a reference point as the only Australian region in which *L. sativae* is presently established.

**Figure 2.**
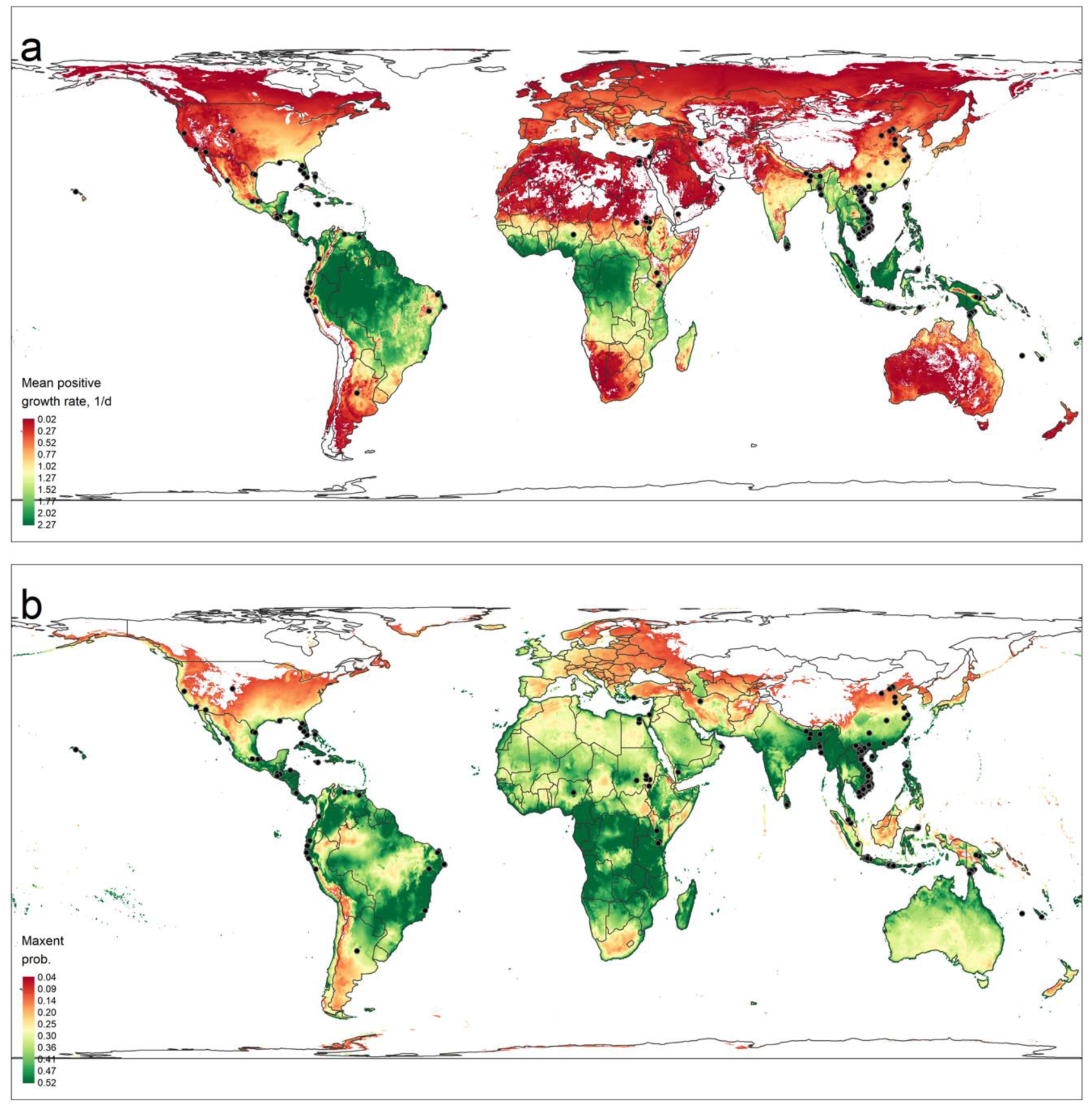
Predicted distribution of *Liriomyza sativae* using a ‘bottom-up’ population growth model (a) and a ‘top-down’ model based on inference from known historical occurrences (b).

**Figure 3.**
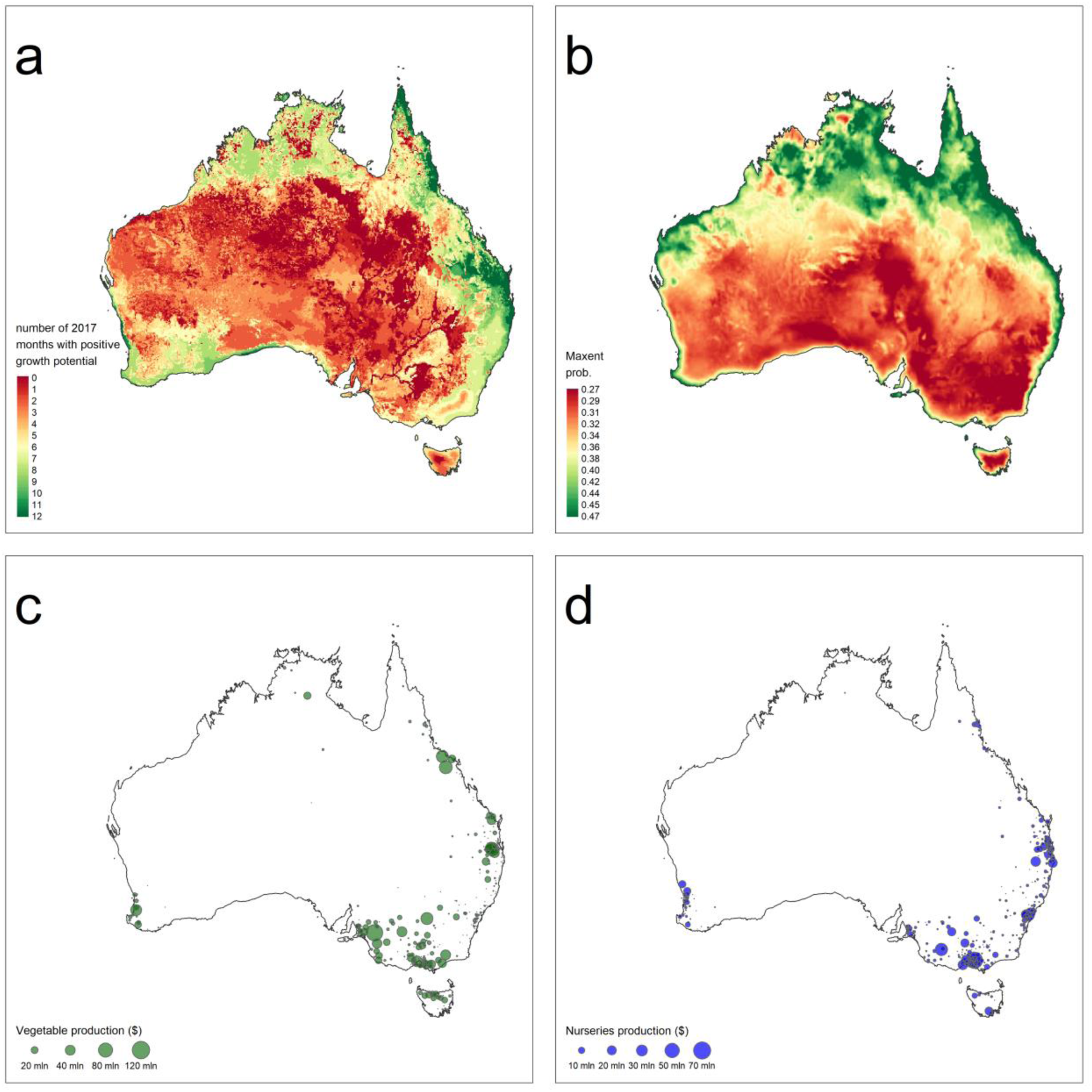
Predicted potential Australian distribution of *Liriomyza sativae* as measured by active months using a ‘bottom-up’ population growth potential model based on temperature and moisture constraints (a) and predicted occurrence probability using a ‘top-down’ MaxEnt model based on inference on known historical occurrences and environmental covariates (b). The local value of vegetable production in 2016 (excluding mushrooms) (c) and nurseries in 2016 (d) as measured by the Australian Bureau of Statistics (2018) are plotted at the centroid of Australian statistical divisions (SA2) where the size of circles represents value of production.

**Figure 4.**
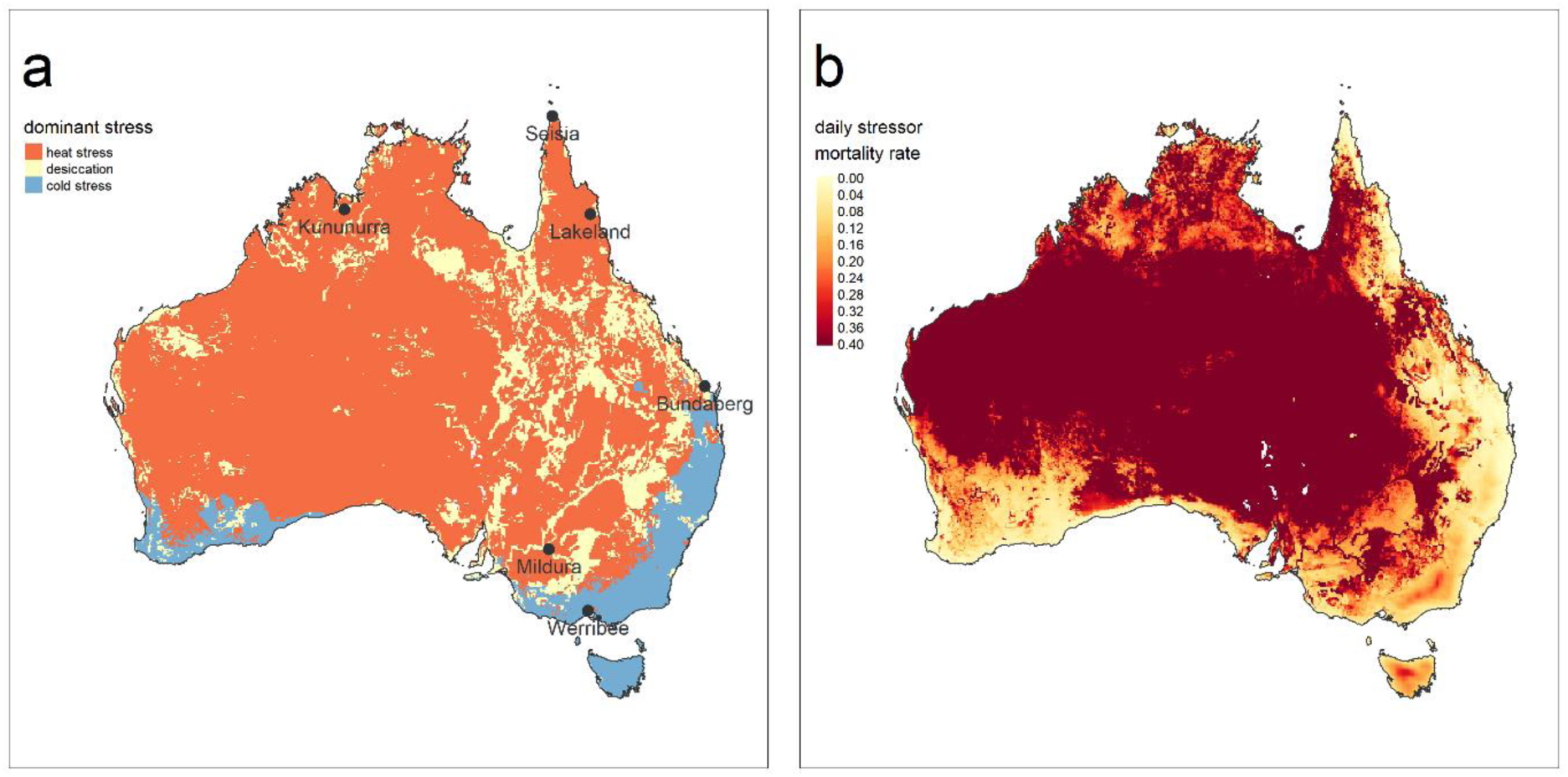
Dominant stressors of *Liriomyza sativae* across Australia based on the highest mortality rate from desiccation, cold, and heat stress across the year (a). The mean daily mortality rate from all stressors was estimated across 2018 (b) with the legend range truncated to emphasise variation in climatic stress across key agricultural regions.

**Figure 5.**
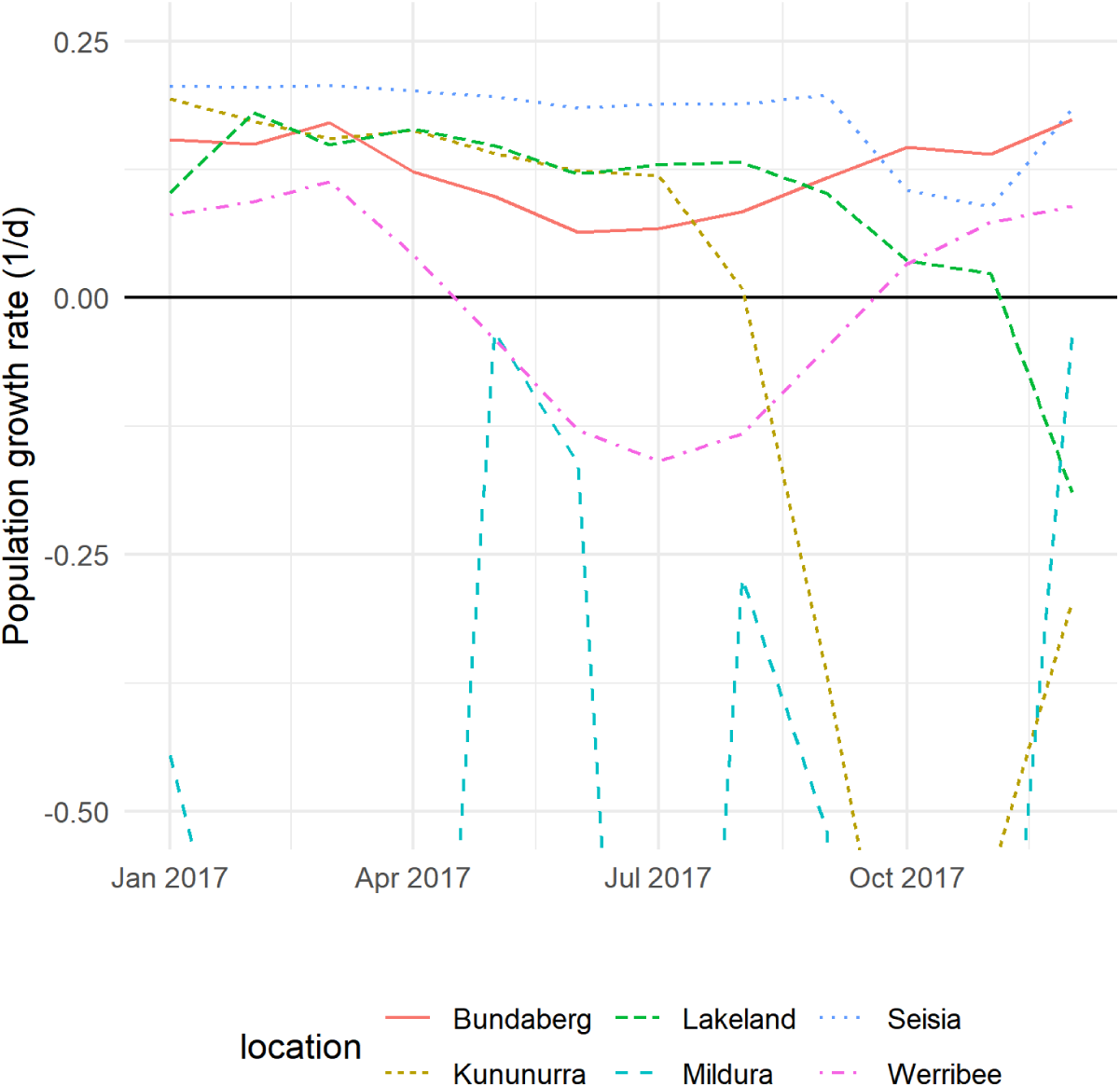
Daily population growth rate of *Liriomyza sativae* associated through time for each month at six agriculturally significant and climatically distinct regions. Locations are shown in Figure 4a.

## 3. Results

### Global distribution of *L. sativae*

In total, 336 unique occurrence records were compiled from published studies spanning 36 countries (see Supplementary Information). The ‘bottom-up’ population growth model captured the known current global distribution of *L. sativae* (Figure 2), predicting strong latitudinal trends in suitability, with more tropical climates with low temperature and moisture stress generally being more favourable. Large areas currently unoccupied by *L. sativae* were predicted to support positive population growth during a part or all of the year, including, India, Europe and southern countries of the African continent, suggesting this species has the potential to significantly expand its current global range. The model had difficulty in predicting occurrence locations in Egypt and Yemen.

### Australian climatic suitability

Large regions of Australia were predicted to have climates capable of supporting positive population growth for *L. sativae* (Figure 3). In particular, large areas in the north-east coastal region were predicted to support positive population growth throughout the year, while northern Australia possessed large regions suitable for approximately half the year. Less suitable were the Mediterranean climatic regions of Australia (e.g. south-west) and temperate regions (e.g. south-east), which were typically predicted to be suitable between one and six months of the year. The arid deserts of central Australia were predicted to be unsuitable for all months of the year.

The explicit inclusion of stress associated mortality facilitated prediction of the dominant stressors limiting population growth in Australia (Figure 4). Regions of lowest mean stress coincided with the far north of Queensland; the region most proximate to the invasion front of *L. sativae*. Cold-stress was the dominant cause of mortality in the temperate south-east, and to a lesser extent, some parts of the south-west. In nearly all other regions of Australia, heat or desiccation were the dominant stressors. This was particularly true for regions furthest away from the coast.

### Vegetable and nursery production

An overview of the vulnerability of affected production zones was provided by overlaying the local production value of vegetable and nursery commodities (Figure 3). Except for northern Victoria, which was predicted to be unsuitable for *L. sativae* due to high heat and desiccation stress (Figure 4), all significant areas of vegetable or nursery production were suitable for *L. sativae* for at least part of the year. Simultaneously, there are large differences in the suitability of these different regions in terms of the number of months per year estimated to support positive population growth. The significant vegetable production regions surrounding the north-eastern coast (e.g. Cairns), melon production regions in the Northern tropics (e.g. Katherine), and nurseries proximate to the eastern coast (e.g. Brisbane) of Australia were frequently predicted to be suitable for *L. sativae* for over half of the year, while vegetable production in southern regions were predicted to be at risk for only part of the year.

The stress map (Figure 4) also suggests how human augmentation of climatic conditions is likely to most exacerbate risks of population outbreaks. In cold limited southern Australia, use of grow houses designed to raise temperatures of crop environments would be expected to increase the number of suitable months. Similarly, in water stressed regions of eastern Australia, irrigation of crops will have the largest predicted impact on population growth potential.

Analysing the impacts of seasonality across a range of key production regions revealed strong monthly variation in population growth rates (Figure 5). In tropical production regions of Lakeland and Kununurra, the dominant stressor is heat and positive growth is achievable most of the year, however with notable population declines predicted during the hot summers. In temperate Bundaberg, positive growth is achievable all year long, with only some deceleration in the winter as a result of cooler temperatures. In temperate Werribee, growth rates are slower throughout the year, and growth becomes negative during winter. Populations are estimated to be able to grow roughly half the year. Finally, the more arid Mildura, VIC, is predicted to be highly unsuitable, with a combination of heat and desiccation stress in the summer, and cold and desiccation stress during the winter, preventing positive growth at any time of the year.

### ‘Bottom up’ versus ‘top down’

Like the population growth model, the ‘top-down’ MaxEnt model tended to predict tropical regions as more favourable for *L. sativae*. However, the variables underpinning this prediction were quite different (Figure 6). Of the four covariates with the highest influence in the MaxEnt model, two of these related to temperature variability; one at the daily scale (BIO 2) and the other at the yearly scale (BIO 7), with both negatively associated with suitability. The two remaining variables of highest influence were related to cold stress: ‘minimum temperature of the coldest month’ (BIO 6), and ‘mean temperature of coldest quarter’ (BIO 11). In contrast, cold stress was predicted as the least important stressor in the population growth model (Figure 1). As might be expected in any model extrapolation exercise, there was generally greater divergence in predictions for areas without occurrence records, including countries with colder climates, such as Russia and Canada.

**Figure 6.**
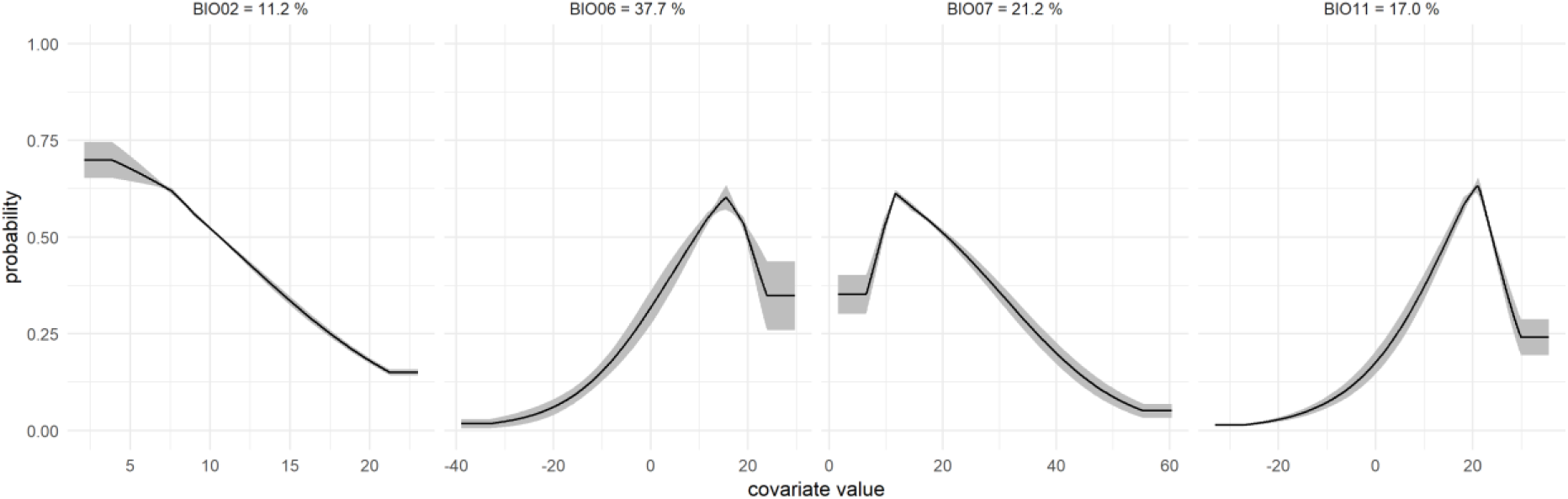
Separated dependence of MaxEnt predicted suitability for the four covariates with the highest percentage model contribution as measured by summed regularised gain (relative influence shown in panel titles), which included ‘mean diurnal range’ (BIO 2), ‘minimum temperature of the coldest month’ (BIO 6), ‘temperature annual range’ (BIO 7), and ‘mean temperature of coldest quarter’ (BIO 11). Lines show the mean of replicates with standard errors shown in grey.

Despite these differences, both models predicted the high suitability of north-eastern Australia (Figure 3), and the low suitability of Australia’s arid interior. While there were some discrepancies in relative suitability, including coastal regions of western Australia, there was generally high alignment in the relative suitability of Australia in both approaches, which strengthens the notion that there are significant areas of Australia presently unoccupied by *L. sativae* that are likely to be climatically suitable and support new populations as its range expands.

## 4. Discussion

The global distribution of *L. sativae* was successfully modelled using two fundamentally different methods: a ‘bottom-up’ approach that explicitly considered determinants of population growth from first principles and biological knowledge, and a ‘top-down’ approach that inferred associations with environmental co-variates using the MaxEnt algorithm. These two contrasting approaches were taken to leverage the relative strength of ‘bottom-up’ models in extrapolating estimates into novel environmental conditions (where processes still apply, but not necessarily correlations), and of ‘top-down’ models to identify latent and complex relationships in high-dimensional ecological variate space (Dormann et al. 2012). These complementary approaches each converged on the conclusion that there are significant areas presently unoccupied by *L. sativae* with high predicted ecological suitability in Australia. The European and Mediterranean Plant Protection Organization (EPPO) has classified *L. sativae* as an A1 pest absent from the EPPO region whereby member countries are recommended to regulate it as a quarantine pest (EPPO 2018). Our findings confirm that *L. sativae* is at risk of establishing in Europe, and supports ongoing prioritisation of biosecurity measures, particularly during warmer months when conditions for population growth are more favourable.

Interestingly, ecological variables with the highest influence that emerged after fitting the MaxEnt model did not align with processes included in the population growth model. While ‘minimum temperature of the coldest month’ (BIO 6), and ‘mean temperature of coldest quarter’ (BIO 11) were found to be highly influential in the ‘top-down’ model, compiled physiological data on the cessation of development and cold stress mortality used to parameterise the ‘bottom-up’ model suggests *L. sativae* has some potential for cold tolerance, with significant adult mortality occurring only after 0°C (Oatman and Michelbacher 1959). This likely explains some of the discrepancies between model predictions into Canada and Russia, with summer months predicted to support population growth in the ‘top down’ model but not the ‘bottom up’ model. To better parameterise this process, more information on cold stress, including effects on the various life-stages (i.e. egg, larva, pupa and adult), is required. Likewise, temperature stability (both daily and annual) was identified as an influential predictor in the MaxEnt model, but fluctuating temperatures were not assumed to directly affect population growth (i.e. in the ‘bottom-up’ model, predicted growth was constant across different fluctuating temperature regimes so long as the total duration at each temperature across regimes was constant). The null expectation was assumed due to insufficient data on *L. sativae*, as negative, positive, and neutral impacts of fluctuating temperature regimes on development have all been observed in different arthropods (Colinet et al. 2015).

In Australia, *L. sativae* is classed under the Emergency Plant Pest Response Deed (EPPRD) as Category 3 (moderate public impact), based on the anticipated negative impact on agricultural industries, and additional public costs (e.g. damage to amenities, or trade restriction). Here we confirmed the expected distribution of *L. sativae* presents a serious risk to Australia’s significant vegetable production and nursery industries. In particular, melon production regions of the Northern Territory, as well as vegetable production and nurseries across coastal regions of Australia’s north-east were identified as having high suitability. However, within suitable locations there will be marked differences in the seasonal pressure of *L. sativae*. Higher annual suitability is associated with higher exposure to incursion events more likely to lead to establishment (i.e. when conditions during incursion are favourable). In contrast, areas containing significant production regions in southern Australia were predicted to support positive population growth for relatively fewer months of the year. This smaller window of suitability implies that these regions will be less vulnerable to incursion events. However, once an exotic pest becomes widely established in a region, availability of micro-climates (e.g. shaded creek beds versus open fields) will likely buffer some local populations against stressful weather conditions (Kearney and Porter 2009). Moreover, establishment of *L. sativae* in areas with year-long suitability is expected to increase the risk of further incursions, or seasonal dispersal events, into areas with partial year suitability (Mitchell et al. 1991). While growing regions in northern Victoria were predicted as highly unsuitable due to heat and moisture stress, irrigation that is known to occur throughout the area will likely create local areas of suitability (as has occurred previously in the American states of Arizona and California). This highlights the importance of domestic biosecurity in such areas where the pest is highly unlikely to spread via natural means.

The simulations undertaken here consider high level climatic factors to determine geographic and seasonal suitability for exotic pest populations, however smaller scale factors will also play a role in determining the realised range of a pest post incursion. A successful incursion requires not just a suitable climate, but the availability of preferred hosts. For example, the red-banded mango caterpillar (*Deanolis sublimbalis*) has remained in the far north of Queensland for nearly two decades, due likely to the lack of mango hosts (Royer 2009). For agricultural pests, a mismatch between the highest risk periods for establishment potential and the seasonality of cultivated crops could theoretically provide a barrier to pest establishment. However, in the case of *L. sativae*, the most suitable climates for *L. sativae* establishment often overlap with production periods for high risk crops (HIA 2019). While cultivated host availability is unlikely to be a problem for *L. sativae* within production regions, the availability of wild hosts will determine the pest’s ability to disperse naturally between production regions, and to remain present in a production region post harvest.

*L. sativae* attacks a very wide range of non-cultivated plants (Oatman and Michelbacher 1959; Spencer 1973), including exotic weeds that are spread widely across Australia and are often associated with disturbed areas such as ports and campgrounds. For example, during surveillance work conducted during 2018 and 2019, one of the most highly preferred weed hosts, siratro (*Macroptilium atropurpureum)*, was observed to be present growing on the fence lines of every shipping port visited, including those at Weipa, Seisia, Thursday Island, Horn Island, and Cairns, Queensland, and was also common within urban areas (Elia Pirtle, pers. obs.). Siratro is present across Australia, as is castor bean (*Ricinus communis*), another highly preferred weed host of *L. sativae* (Atlas of Living Australia website at http://www.ala.org.au. Accessed 29 October 2018). Despite the current isolation of *L. sativae*, it seems unlikely that it will be significantly restricted by host plant availability between production regions, should it make its way further south. The success of invading populations may also be influenced by existing communities of species, both through competitive interactions, and biological control. In Australia, there already exists a considerable, but poorly documented community of agromyzid leafminer species (e.g. *L. brassicae)*, some of which show notable overlap with the preferred host plants of *L. sativae* (Spencer 1973, 1977). Moreover, these agromyzid species are known to support at least 34 genera of parasitoid wasps in Australia, many of which attack *L. sativae* overseas (Ridland 2019, *in review*). Although not considered in the models, Australian parasitoid wasp communities will represent another factor in mitigating establishment and will form the focus of industry communications with an aim to minimise disruption from inappropriate chemical applications.

Although this study is the first to estimate the global distribution of *L. sativae* via a ‘top down’ approach with distributional data, there is unpublished work of note. Using CLIMEX software, Jovicich (2009) modelled the potential future geographic distribution in Australia for four key *Liriomyza spp*. CLIMEX is an intermediary approach between ‘bottom-up’ and ‘top-down’ approaches and has been widely used in forecasting the establishment potential of exotic species through relating development time to temperature (Sutherst and Maywald 1991). However, biological interpretation, theoretical coherence, and parameter estimation is hampered by the use of indices throughout (e.g. the suitability index which is the product of several other indices). In CLIMEX models, taking the product of dimensionless indices contrasts with the ‘bottom up’ model presented here, where all quantities used maintain their physical dimension. This difficulty of interpreting CLIMEX indices also causes difficulties in estimating parameters, which are adjusted manually until predictions align with the observed distribution. There has also been some limited research using ‘bottom up’ approaches to estimate *L. sativae* distributions. Mujica et al. (2014) measured life history data on *L. sativae* at different temperatures and estimated development rates globally using temperature. However, only considering temperature (moisture was not considered) and ignoring mortality at extreme conditions outside the range of positive population growth) this approach predicted very high suitability in arid regions, such as the Saharan desert and central Australia, which we consider to be an erroneous prediction.

With the increasing availability and ease of developing species distribution modelling methods for invading species there has been an accelerating application of such approaches to biological invasions (Jiménez-Valverde et al. 2011). This is generally conducted with a view to forecast establishment risk so that biosecurity resources can be directed with maximum effect. Here we leveraged both strengths of correlative ‘top-down’ and mechanistic ‘bottom-up’ models to estimate for the first time the establishment potential of a recent exotic pest arrival to Australia. Our findings will assist in the development pro-active management, industry preparedness, and biosecurity policies for *L. sativae* in Australia.

## Author contributions

All authors contributed to the study conception and design. Data collection was conducted by James Maino and Peter Ridland. Analysis were performed by James Maino. The first draft of the manuscript was written by James Maino and all authors commented on previous versions of the manuscript. All authors read and approved the final manuscript.

## Acknowledgements

This research was supported by funding from Hort Innovation as part of the RD&E program for control, eradication and preparedness for vegetable leafminer (MT16004).

## Compliance with Ethical Standards

The authors declare no conflict of interest

